# Imaging meningeal inflammation in CNS autoimmunity identifies a therapeutic role for BTK inhibition

**DOI:** 10.1101/2020.08.26.268250

**Authors:** Pavan Bhargava, Sol Kim, Arthur Anthony Reyes, Roland Grenningloh, Ursula Boschert, Martina Absinta, Carlos Pardo-Villamizar, Peter Van Zijl, Jiangyang Zhang, Peter A Calabresi

## Abstract

Leptomeningeal inflammation in multiple sclerosis is associated with worse clinical outcomes and greater cortical pathology. Despite progress in identification of this process in multiple sclerosis patients using post-contrast FLAIR imaging, early trials attempting to target meningeal inflammation have been unsuccessful. There is a lack of appropriate model systems to screen potential therapeutic agents targeting meningeal inflammation. We utilized ultra-high field (11.7 Tesla) MRI to perform post-contrast FLAIR imaging in SJL/J mice with experimental autoimmune encephalomyelitis induced using immunization with proteolipid protein peptide (PLP_139-151_) and complete Freund’s adjuvant. Imaging was performed in both a cross-sectional and longitudinal fashion at time-points ranging from 2 to 14 weeks post-immunization. Following imaging, we euthanized animals and collected tissue for pathological evaluation, which identified dense cellular infiltrates corresponding to areas of contrast-enhancement involving the leptomeninges. These areas of meningeal inflammation contained B cells (B220+), T cells (CD3+) and myeloid cells (Mac2+). We also noted features consistent with tertiary lymphoid tissue within these areas – presence of peripheral node addressin (PNAd) positive structures, CXCL13 producing cells and FDC-M1+ follicular dendritic cells. In the cortex adjacent to areas of meningeal inflammation we identified astrocytosis, microgliosis, demyelination and evidence of axonal stress/damage. Since areas of meningeal contrast enhancement persisted over several weeks in longitudinal experiments, we utilized this model to test the effects of a therapeutic intervention on established meningeal inflammation. We randomized mice with evidence of meningeal contrast enhancement on MRI scans performed at 6 weeks post-immunization, to treatment with either vehicle or evobrutinib (a Bruton’s tyrosine kinase inhibitor) for a period of 4 weeks. These mice underwent serial imaging and we examined the effect of treatment on the areas of meningeal contrast enhancement and noted a significant reduction in the evobrutinib group compared to vehicle (30% reduction versus 5% increase; *P* = 0.003). We utilized ultra-high field MRI imaging to identify areas of meningeal inflammation and to track them over time in SJL/J mice with experimental autoimmune encephalomyelitis and then utilized this model to identify Bruton’s tyrosine kinase inhibition as a novel therapeutic approach to target meningeal inflammation. The results of this study provide support for future studies in multiple sclerosis patients with imaging evidence of meningeal inflammation.

## Introduction

Multiple sclerosis is a demyelinating disorder of the CNS that has both inflammatory and demyelinating components (Jones *et al*., 2009). Leptomeningeal inflammation was first noted in patients with secondary progressive multiple sclerosis in 2004 (Serafini *et al*., 2004) and subsequently has been described in primary progressive multiple sclerosis and relapsing-remitting multiple sclerosis (Lucchinetti *et al*., 2011; Choi *et al*., 2012). Meningeal inflammation may be organized as ectopic lymphoid follicles consisting of B cells, follicular-helper T cells and follicular dendritic cells or may be unorganized consisting of lymphocytic and monocytic populations (Magliozzi *et al*., 2007; Absinta *et al*., 2015). The presence of meningeal inflammation is linked to greater cortical demyelination and a more severe disease course, suggesting a role for meningeal inflammation in multiple sclerosis disease progression (Magliozzi *et al*., 2007; Howell *et al*., 2011). In people with multiple sclerosis contrast-enhanced FLAIR imaging appear to detect areas of leptomeningeal contrast enhancement that correspond to areas of meningeal inflammation on autopsy (Absinta *et al*., 2015). More recent studies with 7 Tesla MRI appear to demonstrate enhanced ability to detect this process in multiple sclerosis patients (Harrison *et al*., 2017; Zurawski *et al*., 2019). There is however still controversy regarding the best approach to identify this phenomenon in multiple sclerosis patients and its relationship to cortical damage, however all studies have shown a consistent relationship between cortical thinning and the presence of leptomeningeal enhancement (Absinta *et al*., 2015, 2017; Harrison *et al*., 2017; Makshakov *et al*., 2017; Zivadinov *et al*., 2017; Bergsland *et al*., 2019; Zurawski *et al*., 2019; Absinta and Ontaneda, 2020). Trials have been conducted to target meningeal inflammation (Komori *et al*., 2016; Topping *et al*., 2016; Bergman *et al*., 2018; Bhargava *et al*., 2019), but these attempts are hampered by the lack of appropriate models to screen potential therapies especially those that would impact established meningeal inflammation.

Meningeal inflammation has been noted in animal models of multiple sclerosis, including the relapsing remitting model of experimental autoimmune encephalomyelitis (rr-EAE) in SJL/J mice (Magliozzi *et al*., 2004; Pikor *et al*., 2015; Ward *et al*., 2019). While meningeal inflammation is apparent in EAE prior to the onset of inflammation in the brain parenchyma, the cellular composition and contributors at this stage differ from those seen in multiple sclerosis meningeal tissue (Christy *et al*., 2013; Parker Harp *et al*., 2019). In experimental autoimmune encephalomyelitis (EAE) models with chronic expression of the B cell chemokine CXCL-13 and BAFF (B-cell activating factor) in the CNS, evidence of meningeal inflammation with an abundance of B cells and evidence of tertiary lymphoid organ formation has been described (Serafini *et al*., 2004). In the rr-EAE model, the occurrence of leptomeningeal inflammation is variable and the composition of meningeal infiltrates appears to evolve over time – from predominantly T cell to a T and B cell infiltrate but has not been systematically studied (Pikor *et al*., 2015). There is also evidence for increased demyelination and axonal damage in areas of the spinal cord or cortex adjacent to areas of leptomeningeal inflammation in EAE similar to that seen in multiple sclerosis (Pikor *et al*., 2015; Ward *et al*., 2019). Thus, the leptomeningeal inflammation noted in rr-EAE mice appears to recapitulate several features seen in multiple sclerosis. However, since leptomeningeal inflammation is not uniform between mice, studying the effect of therapies on this phenomenon is challenging. Identification of meningeal inflammation on imaging and tracking its evolution over time provides an outcome that could be used to study the effect of interventions targeting meningeal inflammation. In this study we first describe the application of high-field contrast-enhanced MRI to identify and track meningeal inflammation in the rr-EAE model in SJL/J mice, providing a novel approach to screen therapies that could target this pathophysiological process.

A rational target potentially involved in leptomeningeal inflammation is Bruton’s tyrosine kinase (BTK), transmitting signals through a variety of receptors leading to B cell and myeloid cell activation and pro-inflammatory polarization (Gabhann *et al*., 2014; Haselmayer *et al*., 2019). From the various BTK inhibitors developed for clinical use, Evobrutinib was the first one to show clinical efficacy in EAE and in a relapsing-remitting multiple sclerosis Phase 2 clinical trial (Montalban *et al*., 2019). We here utilize this model to identify the therapeutic effects of Evobrutinib on meningeal inflammation, providing evidence for potential utility of this class of medications in targeting this pathological process in patients with multiple sclerosis.

## Methods

### Animals

SJL/J mice were purchased from Jackson Laboratories for all experiments. All mice were maintained in a federally approved animal facility at the Johns Hopkins University in accordance with the Institutional Animal Care and Use Committee. Female mice 7–8 weeks of age were used in all of the experiments and were housed in the animal facility for one week prior to start of experiments.

### Induction of relapsing remitting EAE

Female SJL/J mice were immunized subcutaneously at two sites over the lateral abdomen with 100 ug of PLP139-151 peptide with complete Freund’s adjuvant (CFA) containing 4 ug/ml Mycobacterium tuberculosis H37RA (Difco laboratories). Mice were weighed and scored serially to document disease course. Scoring was performed using the following scale: 0-normal, 1-limp tail, 2-hind limb weakness, 3-hind limb paralysis, 4-fore limb weakness, 5-death.

### Imaging of meningeal disease

At the time points noted above, a horizontal 11.7 Tesla Bruker scanner (Bruker Biospin, Billerica, MA) with a triple-axis gradient system (maximum gradient strength = 740 mT/m). A 72 mm volume transmit coil and 4-channel receive-only phased array coil was used to image the mouse forebrain, whereas the mouse cerebellum was imaged with a quadrature surface transmit/receive cryogenic probe to achieve high resolution. During imaging, mice were anesthetized with isoflurane together with air and oxygen mixed (3:1 ratio) and respiration was monitored via a pressure sensor and maintained at 60 breaths/min. Before imaging, 0.1 mL diluted Magnevist (gadopentetate dimeglumine, Bayer HealthCare LLC, Whippany, NJ, 1:10 with PBS) was injected I.P. T2-weighted images were acquired using a rapid acquisition with refocused echoes (RARE) sequence with the parameters: echo time (TE)/repetition time (TR) = 30/5000 ms; echo train length = 8, field of view = 15 mm x 15 mm, 30 axial slices with a slice thickness of 0.5 mm to cover the entire brain, matrix size = 192 × 192, and a signal average of 4. T1-weighted images were acquired using a spin echo sequence with the same parameters as the T2-weighted images except TE/TR = 9/300 ms, matrix size = 128 × 128, and only 15 axial slices were acquired to cover the fore-brain region. FLAIR images were acquired using the RARE sequence with an inversion pulse. The imaging parameters were: TE/TR = 20/3000 ms, steady-state inversion time = 1000 ms, 15 axial slices with slice thickness of 0.5 mm, the same matrix size and imaging geometry as the T1-weighted images. Scans were then analyzed to identify areas of meningeal contrast enhancement by two independent examiners.

We utilized a semi-quantitative scheme to quantify the extent of meningeal contrast enhancement – we counted the number of areas of meningeal contrast enhancement on each individual MRI slice and used the cumulative number as a representation of the amount of meningeal contrast enhancement. All quantifications were performed by at least two independent examiners and their scores were averaged.

### Histological identification of meningeal disease

Following the final imaging time point, mice were deeply anesthetized with sodium pentoparbital (100 mg/kg) followed by intra-cardiac perfusion with 4% paraformaldehyde. The skull was separated, stripped of all soft tissue and placed in 14% EDTA at pH 7.6 for decalcification. Skulls were weighed daily to determine the time point of maximal decalcification. The decalcified skulls were transferred to 30% sucrose for 48 hours and embedded in OCT before snap-freezing. Tissues were cryosectioned at 10-μm thickness onto glass slides (Superfrost Plus, Fisher) to capture the appropriate brain and meningeal areas corresponding to meningeal enhancing lesions noted on MRI. Hematoxylin & Eosin (H&E) staining (Empire Genomics) was performed to verify that the appropriate areas had been captured during this process.

### Immunohistochemistry to characterize meningeal disease

Blocking and permeabilization of sections was performed in PBS containing 5% normal goat serum (NGS) and 0.4% Triton X-100 for 1 h at room temperature followed by incubation overnight at 4°C in PBS containing 3% NGS, 0.1% Triton X-100, and primary antibodies. The optional antigen retrieval was performed with heated citrate buffer (pH 6) for 5–8 minutes before blocking. The slides were then washed and incubated with appropriate secondary antibodies conjugated to Alexa fluorophores (1:1000, Invitrogen), 3% NGS and 0.1% Triton X-100 for 1 h at room temperature. The slides were mounted using anti-fade reagent with DAPI (Prolong Gold Anti-fade, Gibco) and examined under fluorescent microscopes. The antibodies tested included APP, B220, CD3, CXCL13, FDC-M1, Fibrinogen, GFAP, IBA-1, INOS, Mac-2, SMI-32, and peripheral node addressin (PNAd) (See List of Antibodies in Supplementary Table 1). For myelin staining, Luxol Fast Blue (IHC World; IW-3005) was used following the manufacturer’s protocol.

For imaging, Olympus BX41, Axio Cell Observer or ZEISS LSM 800 were used with appropriate exposure and magnification. Each slide was imaged for 1–2 sections from each group (EAE vs Naïve) and each panel was processed with ZEN lite (Zeiss) and National Institutes of Health ImageJ software (NIH, https://imagej.nih.gov/ij/).

### Testing the effect of Evobrutinib treatment on leptomeningeal inflammation

Evobrutinib was obtained from EMD Serono and mice were treated with either vehicle solution (20% kleptose in 100 mM Na-Citrate buffer, pH 3) or evobrutinib dissolved in vehicle solution by daily oral gavage (10 mg/kg) between weeks 6–10 post-immunization. Each group (vehicle vs treatment) underwent contrast-enhanced MRI at week 6, 8, and 10 and brain tissues were collected for histopathology at the final imaging time point. The quantity of meningeal contrast-enhancement was quantified by raters blinded to treatment assignment at each imaging time-point to assess the effect of interventions on meningeal inflammation.

### Statistical analysis

For comparisons of pathological outcomes between groups we used a Mann-Whitney U test. To compare the effects of vehicle versus evobrutinib on meningeal contrast enhancement in mice with EAE we used a mixed effects model since we had repeated measures in each group. All analyses were performed using GraphPad Prism Version 8.3.0.

### Data availability statement

The authors confirm that the data supporting the findings of this study are available within the article and its supplementary material.

## Results

### MRI identifies areas of leptomeningeal contrast enhancement in SJL/J mice with EAE

To determine whether MRI can identify meningeal inflammation in EAE we immunized female SJL/J mice as described in the Methods. The typical course of disease in these mice is depicted in Fig. 1A. We chose the SJL/J rr-EAE model for our experiments since previous studies have reported the development of meningeal inflammation with features of tertiary lymphoid tissue in this model (Magliozzi *et al*., 2004; Pikor *et al*., 2015). In our initial cross-sectional experiments, we performed contrast-enhanced MRI scans at 6 weeks post-immunization. At this time-point, the majority of mice demonstrated one or more contrast-enhancing lesions within the sub-arachnoid space (Fig. 1B). Some of these areas of contrast enhancement were noted on both T1 weighted as well as FLAIR sequences. However, similar to human studies demonstrating the superiority of post-contrast FLAIR imaging to detect leptomeningeal pathology (Tsuchiya *et al*., 2001; Absinta *et al*., 2015), FLAIR imaging revealed more/larger areas of leptomeningeal contrast enhancement (Supplementary Fig. 1). Based on these findings we subsequently utilized FLAIR MRI for detection and monitoring of leptomeningeal contrast enhancement. Leptomeningeal contrast enhancement was detected both over the surface of the cerebral cortex and in the hippocampal fissure (Fig. 1B), but not in control animals (Fig. 1C).

**Figure 1.**
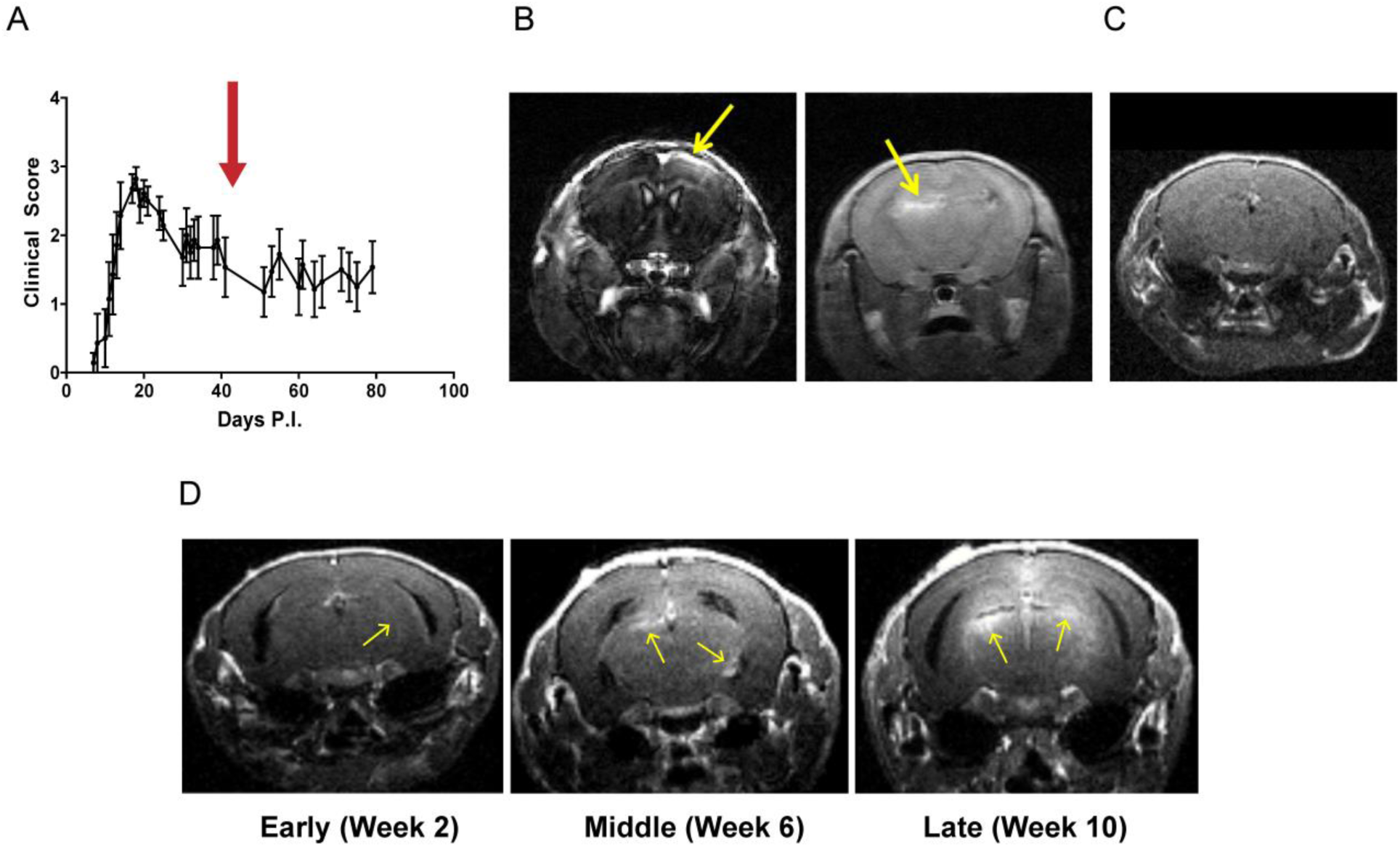
Ultra-high field MRI detects meningeal contrast enhancement in mice with EAE. **(A)** The typical disease course of EAE in SJL/J mice immunized with CFA and PLP131-155. The red arrow corresponds to the time-point of imaging depicted in panel B and C. **(B)** EAE mice at 6 weeks post-immunization underwent MRI following intra-peritoneal administration of gadolinium contrast. Notice the presence of contrast enhancement involving the meninges (yellow arrows) either over the cortical surface (left) or deeper within the hippocampal fissure (right). This was noted both on the MP-RAGE (left) and FLAIR (right) images. **(C)** We did not detect similar contrast enhancement in naïve SJL/J mice of a similar age. **(D)** In subsequent experiments we noted presence of meningeal inflammation as early as two weeks post-immunization and there appeared to be an increase in the frequency and extent of meningeal enhancement over the course of the disease (from 2 weeks to >10 weeks post-immunization).

When performing additional experiments at early (2 weeks post-immunization), middle (4–6 weeks post-immunization) or late (≥8 weeks post-immunization) time points, we found an increase in the number and size of leptomeningeal contrast enhancing lesions at later time points (Fig. 1D).

### Leptomeningeal contrast enhancing areas on MRI correspond to areas of meningeal inflammation

Following completion of imaging, mice were euthanized, and tissues obtained for histopathological evaluation as described in the Methods. We noted the presence of dense cellular infiltrates in the sub-arachnoid space in the regions corresponding to meningeal contrast enhancement seen on MRI (Fig. 2A–D). We then performed immunohistochemistry to identify the cell populations comprising these meningeal infiltrates and found that the major components of the meningeal infiltrates were B220+ B lymphocytes (Fig. 2E and 3A) and CD3+ T lymphocytes (Fig. 2F and 3A). We also identified myeloid cells within these infiltrates expressing Mac-2 (Fig. 3B) or Iba-1 (Fig. 3C) which are likely macrophages.

**Figure 2.**
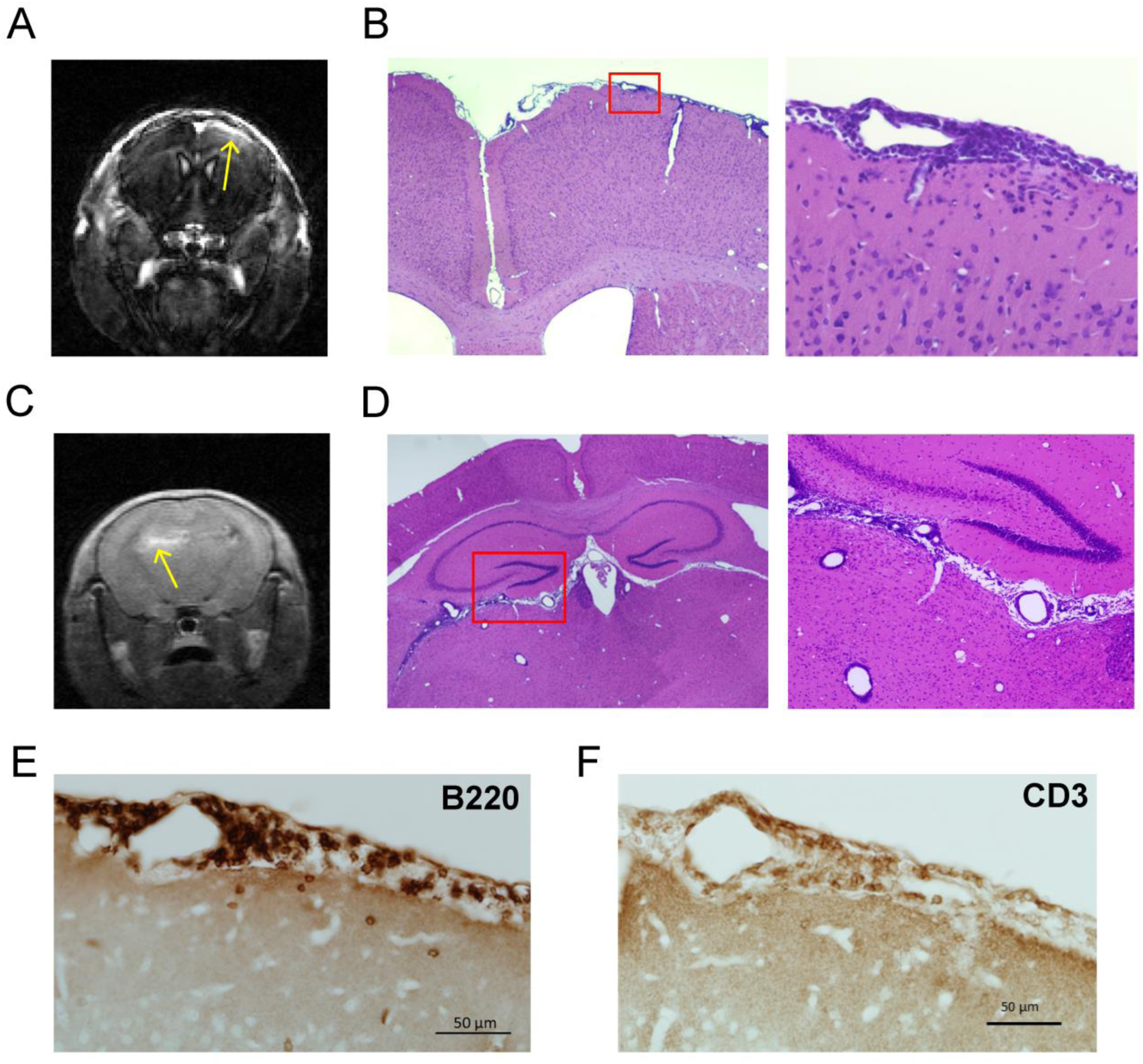
Areas of meningeal contrast enhancement correspond to leptomeningeal inflammation. **(A and C)** Following identification of areas of meningeal contrast enhancement on MRI we euthanized mice and obtained brain tissue as described in the methods section. **(B and D)** On **H and E** staining of the tissue corresponding to MRI areas of meningeal contrast enhancement we noted the presence of dense cellular infiltrates involving the leptomeninges. **(E)** In the area of meningeal contrast enhancement, immunohistochemistry revealed the presence of numerous B220+ B cells and **(F)** CD3+ T cells.

**Figure 3.**
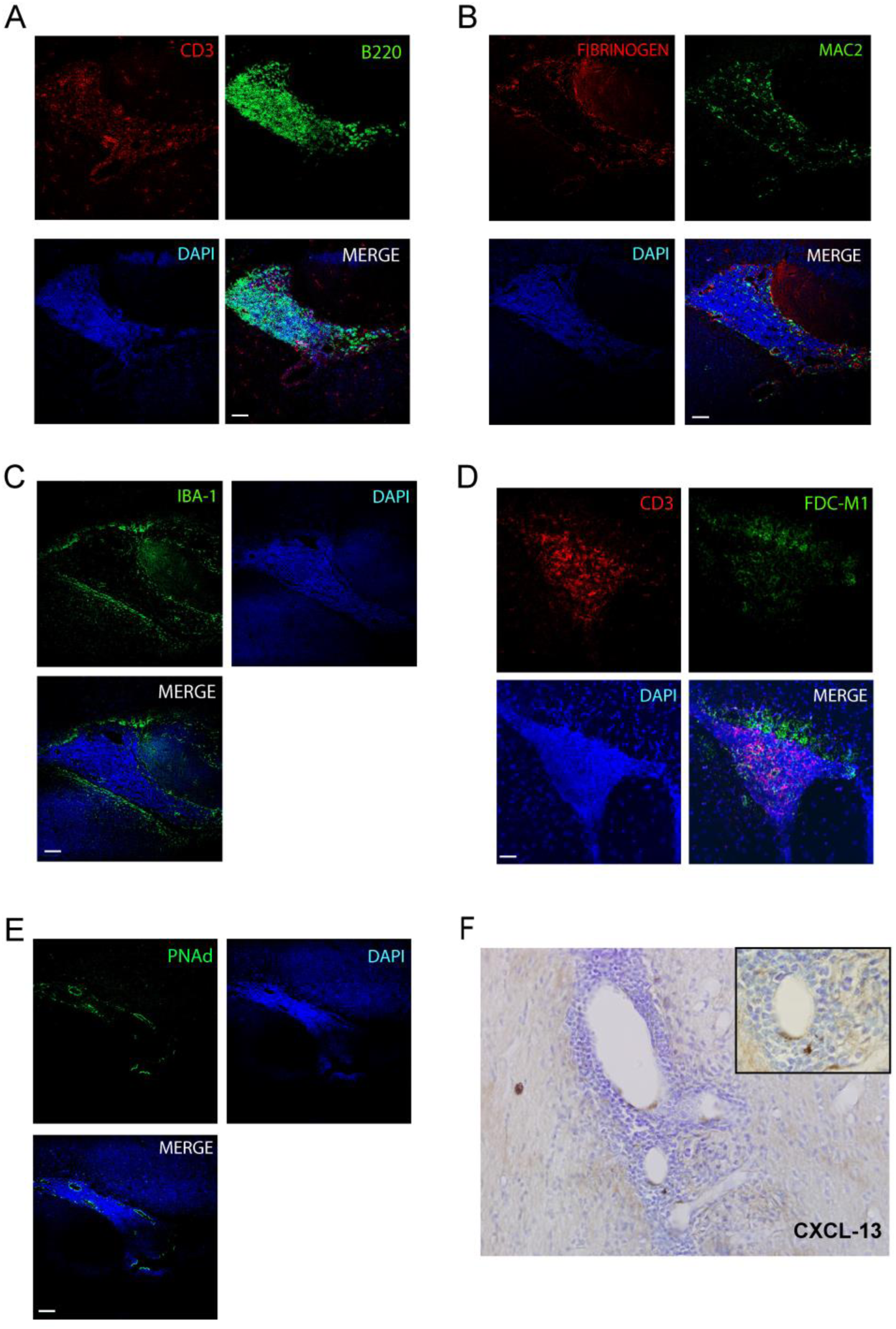
Leptomeningeal inflammatory infiltrates contain lymphocytes, myeloid cells and have markers of tertiary lymphoid tissue. **(A)** Areas of meningeal inflammation demonstrated the presence of numerous CD3+ T cells and B220+ B cells. **(B)** These areas also demonstrated the presence of myeloid cells (Mac2+) and deposition of fibrinogen was noted consistent with breakdown of blood brain barrier resulting in contrast-enhancement. Mac2+ cells appeared to cluster in areas of fibrinogen deposition. **(C)** Iba1+ cells – representing microglia and myeloid cells both within the areas of meningeal inflammation as well as the cortex surrounding this. **(D)** In addition, FDC-M1+ follicular dendritic cells were present within the leptomeningeal inflammatory infiltrates. **(E)** Other features consistent with tertiary lymphoid tissue included the presence of PNAd+ structures that were suggestive of high-endothelial venules and **(F)** CXCL13+ cells within these areas. Scale Bars: 50 µm for all panels.

We noted the presence of blood-meningeal barrier breakdown within these inflammatory infiltrates as evidenced by the presence of fibrinogen deposition (Fig. 3B). Interestingly, the pattern of fibrinogen deposition also appeared to overlap with the distribution of Mac2+ myeloid cells (Fig. 3B). This is not surprising given the recent data regarding the ability of fibrinogen to signal to myeloid cells through the CD11b receptor (Ryu *et al*., 2015).

These results confirm pervious reports of the presence of meningeal inflammation in SJL/J EAE and demonstrate the ability of contrast-enhanced MRI imaging to detect areas of meningeal inflammation in SJL/J mice with EAE.

### Areas of meningeal inflammation display markers of tertiary lymphoid tissue

Since prior reports have noted that meningeal inflammation in a subset of multiple sclerosis patients and mice with EAE demonstrates characteristics of tertiary lymphoid tissue, we performed staining for markers that would help identify this phenomenon. We noted a network of FDC-M1+ follicular dendritic cells within the meningeal inflammatory infiltrates (Fig. 3D). We also stained for PNAd - a marker for high-endothelial venules normally found in lymphoid tissue and identified the presence of PNAd+ vessel like structures within a subset of the areas of meningeal inflammation (Fig. 3E). The presence of follicular dendritic cells and high-endothelial venules within areas of meningeal inflammation suggest that these areas may be organizing to form tertiary lymphoid structures (Luo *et al*., 2019).

In addition to this we noted the presence of cells staining for CXCL-13 (Fig. 3F) a chemokine that is responsible for chemotaxis of B cells and has been identified in ectopic lymphoid tissue (Magliozzi *et al*., 2004).

These results demonstrate the presence of features of tertiary lymphoid tissue in areas of meningeal inflammation in the SJL/J EAE model.

### Meningeal inflammation is associated with pathological changes in the adjacent cortex

Leptomeningeal inflammation has been linked to changes in the adjacent cortex including cortical demyelination and neuronal dropout in patients with multiple sclerosis (Magliozzi *et al*., 2010; Howell *et al*., 2015). We first evaluated the effects on the cortex adjacent to areas of meningeal inflammation by examining changes in astrocytes and microglia. Using staining for GFAP we identified marked astrocytosis in the cortex adjacent to areas of meningeal inflammation (Fig. 4A and C) accompanied by a disruption in the glia limitans.

**Figure 4.**
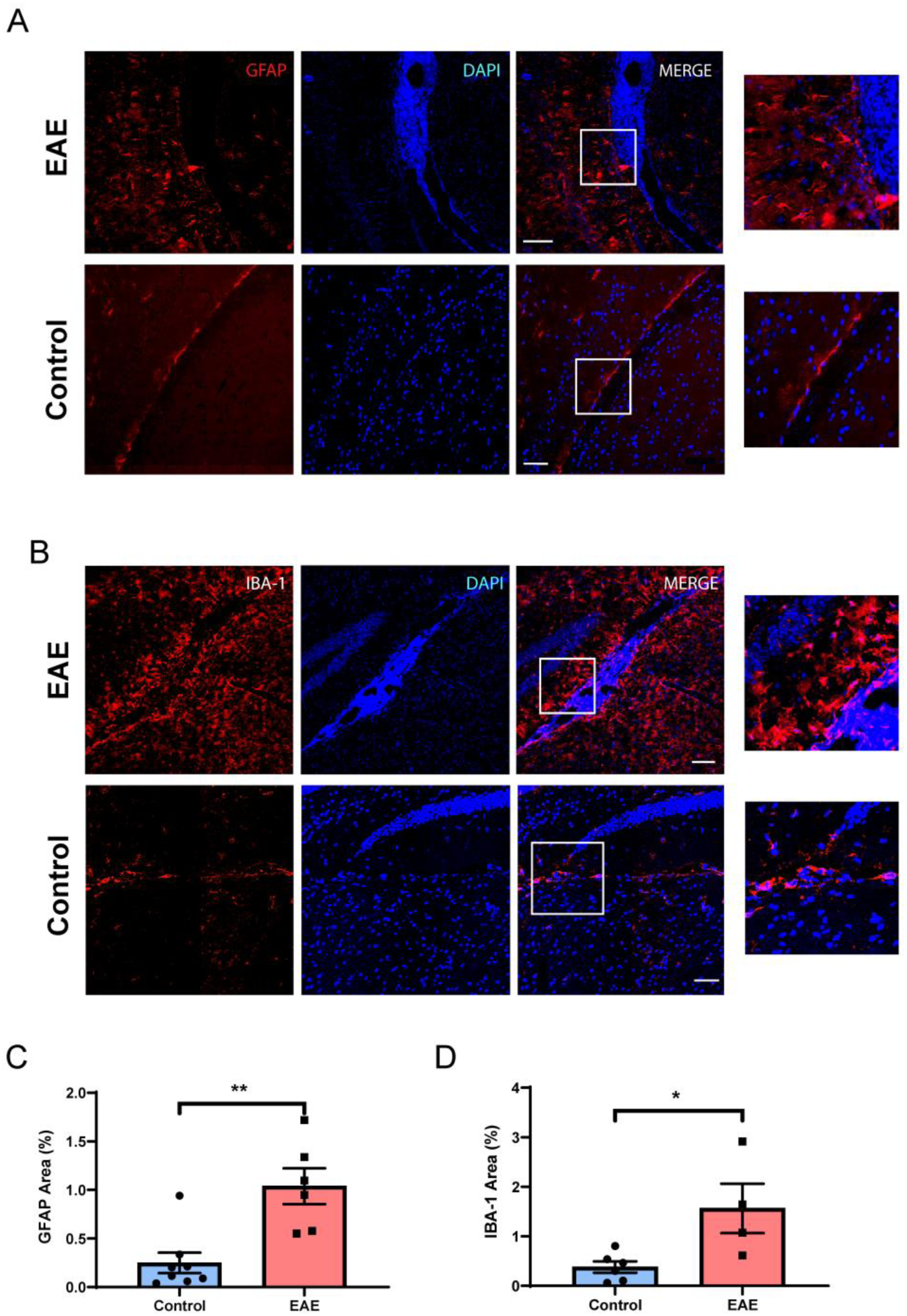
Cortex adjacent to leptomeningeal inflammation demonstrates astrocytosis and microgliosis. **(A)** Areas of the cortex adjacent to meningeal inflammation demonstrated increased numbers of astrocytes as well as a disruption of the glia limitans as compared to control brain. This staining is quantified in **(C). (B)** The cortex adjacent to areas of meningeal inflammation also demonstrated increased numbers of Iba-1 positive cells and these cells appeared more activated compared to control brains. This staining is quantified in **(D)**. Scale bars in **A** and **B** are 100 µm. For **C** and **D** data are shown as mean ± SEM and *P* values are derived from a Mann-Whitney U test and **: *P* < 0.005, *: *P* < 0.05.

In the same areas we also noted an increase in the number of Iba-1+ cells compared to controls (Fig. 4B and D). Iba-1+ cells in the cortex adjacent to meningeal inflammation had an activated appearance as compared to the contralateral hemisphere that was unaffected by meningeal inflammation (Supplementary Fig. 2).

Since recent data have demonstrated that the phenotype of astrocytes can be determined by cross-talk with microglia – and this can lead to the development of reactive neurotoxic astrocytes (Liddelow *et al*., 2017; Rothhammer *et al*., 2018), we examined whether the astrocytes in the cortex adjacent to areas of meningeal inflammation demonstrated markers of this phenotype (C3 and PSMB8). We noted the presence of C3+ and PSMB8+ GFAP+ cells (Supplementary Fig. 3) which potentially represent neurotoxic reactive astrocytes in the cortex adjacent to meningeal infiltrates.

Since leptomeningeal inflammation in multiple sclerosis tissue is linked to greater cortical demyelination and neuronal damage, we evaluated myelin and axonal health in the cortex adjacent to meningeal infiltrates. We performed staining for myelin using Luxol-Fast Blue dye and noted increased vacuolation and decreased staining in cortical areas adjacent to meningeal inflammation compared to controls (Fig. 5A).

**Figure 5.**
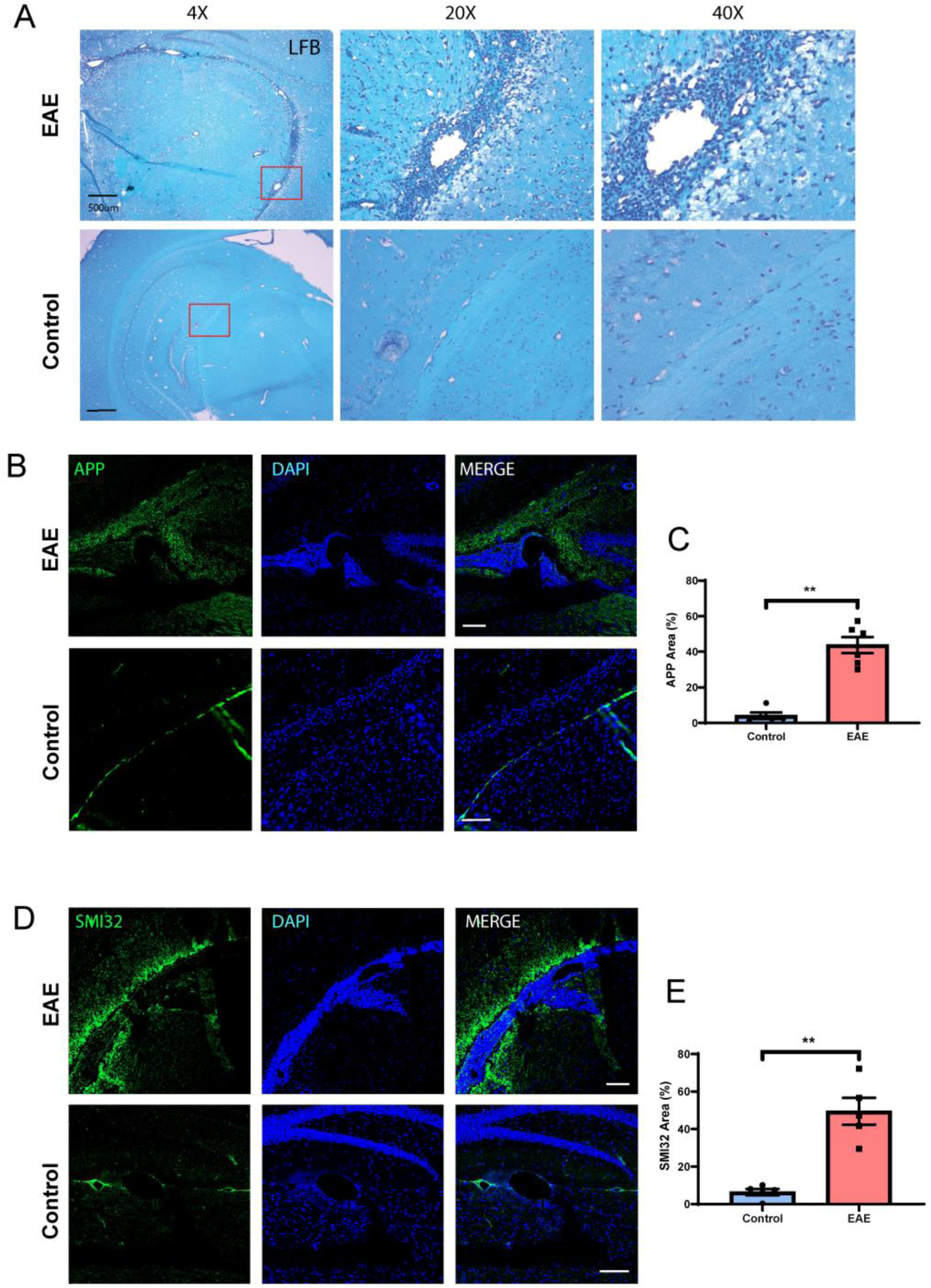
Cortex adjacent to leptomeningeal inflammation demonstrates axonal damage and demyelination. **(A)** We examined myelination using Luxol Fast Blue staining in the cortex adjacent to areas of inflammation and noted increased demyelination (reduced staining and increased vacuolation) compared to normal brains. **(B)** We examined axonal stress/damage using staining for APP and noted increased staining in the cortex adjacent to meningeal inflammation compared to controls. **(C)** Quantification of APP staining in cortex adjacent to areas of meningeal inflammation in EAE compared to similar areas in controls. **(D)** We noted similar increase in staining for non-phosphorylated neurofilament (SMI-32) in the cortex adjacent to areas of meningeal inflammation compared to control brains. **(E)** Quantification of SMI-32 staining in the cortex adjacent to areas of meningeal inflammation in EAE compared to similar areas in controls. Scale bars in **A** are 500 µm, **B** and **D** are 100 µm. Data in **C** and **E** are shown as mean ± SEM and *P* values derived from Mann-Whitney U test with **: *P* < 0.005.

To assess for axonal stress/damage we performed staining for non-phosphorylated form of neurofilament-H (SMI-32) and noted increased staining in the cortex adjacent to meningeal inflammation compared to control tissue (Fig. 5B and C). We also noted similar results using another marker of axonal stress and impaired fast axonal transport – amyloid precursor protein (APP) (Medana and Esiri, 2003) and noted increased staining in the cortex adjacent to areas of meningeal inflammation compared to control tissue (Fig. 5D and E). This is consistent with impaired axonal transport of proteins that can occur with either axonal stress or damage (Schirmer *et al*., 2011).

These results suggest a possible effect of meningeal inflammation on the underlying cerebral cortex with microglial and astrocytic activation, demyelination and axonal damage. This is significant since these findings parallel the demyelination, axonal and neuronal damage noted in progressive multiple sclerosis patients with evidence of meningeal inflammation.

### Longitudinal changes in meningeal enhancement and composition of meningeal infiltrates

Since imaging studies in multiple sclerosis have demonstrated the persistence of areas of leptomeningeal contrast enhancement over time, we performed longitudinal experiments to study the time course of these areas of meningeal inflammation. Mice were imaged serially at early (week 2), middle (week 4 or 6), and late (week 8 or 10) and lesions were identified at each time point. We noted that new lesions formed over time and tended to accumulate over time (Fig. 6A).

**Figure 6.**
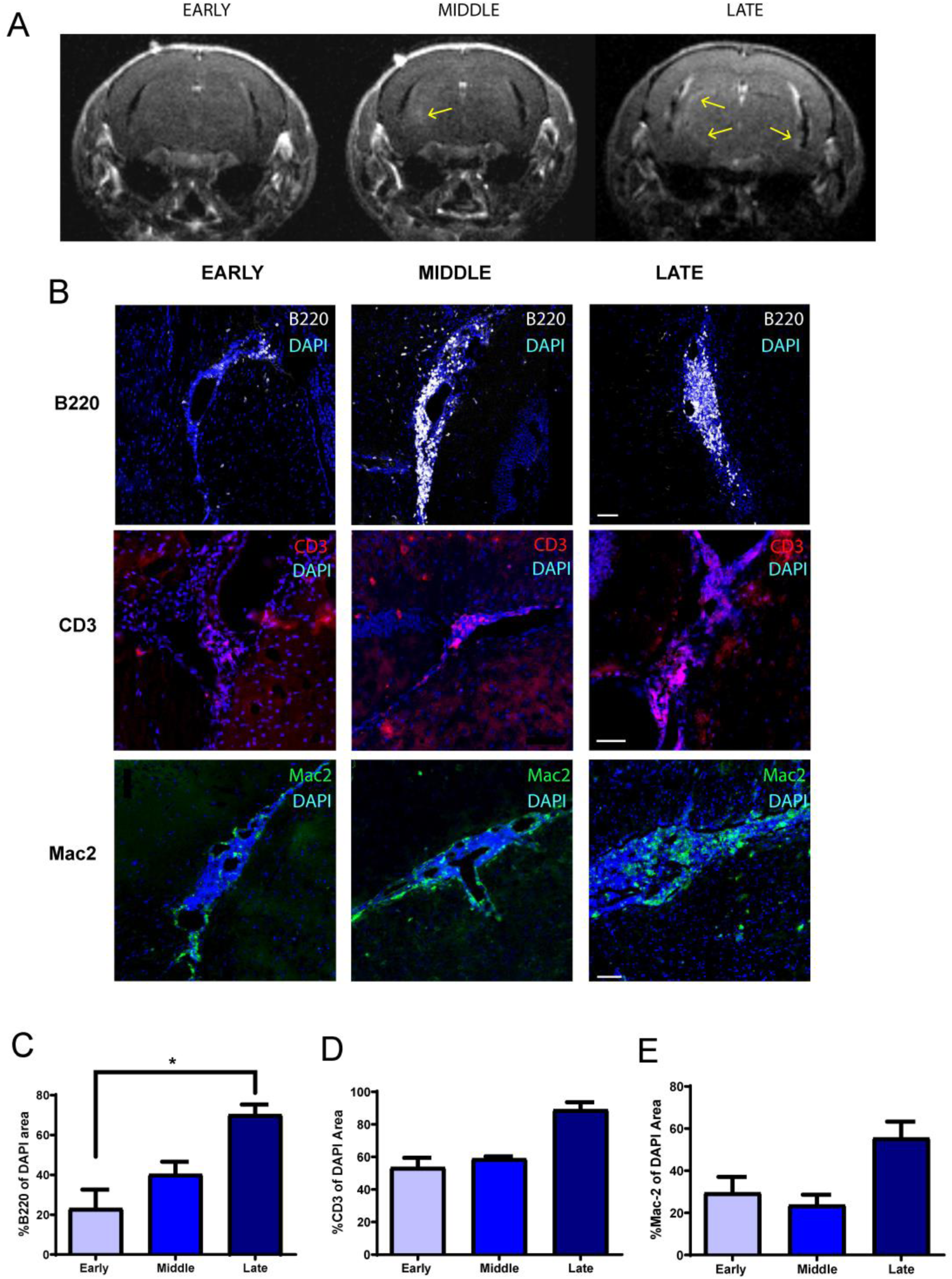
Leptomeningeal contrast enhancement persists over the course of EAE and proportion of B cells increases in areas of leptomeningeal inflammation over time. **(A)** We noted that leptomeningeal contrast enhancement persisted for several weeks and new areas accumulated over the course of EAE. **(B)** We also compared the composition of the meningeal inflammatory infiltrate over the course of EAE and noted changes in immune cell populations with a significant increase in the B cell population within these areas **(C)**. While the T cell **(D)** and myeloid cell **(E)** populations also appeared to increase over the course of EAE – these were not statistically significant. Scale bars in **B** are 50 µm for CD3 panels and 100 µm for B220 and Mac2 panels. Data in **C** and **E** are shown as mean ± SEM and *P* values derived from Mann-Whitney U test with *: *P* < 0.05.

We also examined the change in the composition of meningeal infiltrates over this time period. In these experiments, mice that had evidence of meningeal contrast enhancement on MRI were sacrificed at early, middle or late time points and we performed immunostaining for T cells, B cells and myeloid cells. We noted a steady significant increase in the proportion of B cells in the inflammatory infiltrate over the course of EAE (Fig. 6B and C). This B cell abundance is reminiscent of leptomeningeal inflammation noted in multiple sclerosis patients (Magliozzi *et al*., 2006; Wicken *et al*., 2018). While T cell and myeloid cell numbers also appeared to increase, this change was not statistically significant (Fig. 6B, D and E).

These results demonstrate that meningeal inflammation in rr-EAE can be detected by contrast-enhanced imaging and tends to persist over a period of several weeks providing a potential marker for monitoring response to putative therapies. The composition of meningeal inflammation also evolves over time with a greater B cell abundance noted later in the course of rr-EAE.

### BTK inhibition reduces established meningeal inflammation and leptomeningeal contrast enhancement in SJL/J mice with EAE

Since B cells are a major component of the meningeal infiltrate in multiple sclerosis patients, several trials have unsuccessfully attempted to target meningeal inflammation using anti-CD20 monoclonal antibodies (Svenningsson *et al*., 2015; Komori *et al*., 2016; Topping *et al*., 2016; Bhargava *et al*., 2018). Given the presence of both B cell and myeloid populations in areas of meningeal inflammation, we tested whether an agent that targets both these cell populations would have an effect on leptomeningeal inflammation. We utilized MRI to identify mice with meningeal inflammation and to then evaluate the effect of a BTK inhibitor – evobrutinib, on this process.

Mice with evidence of meningeal contrast enhancement at 6 weeks post-immunization were randomized to either evobrutinib or vehicle (Fig. 7A). We noted a significant reduction in meningeal contrast enhancements in the evobrutinib group but not in the vehicle group (Fig. 7B and C).

**Figure 7.**
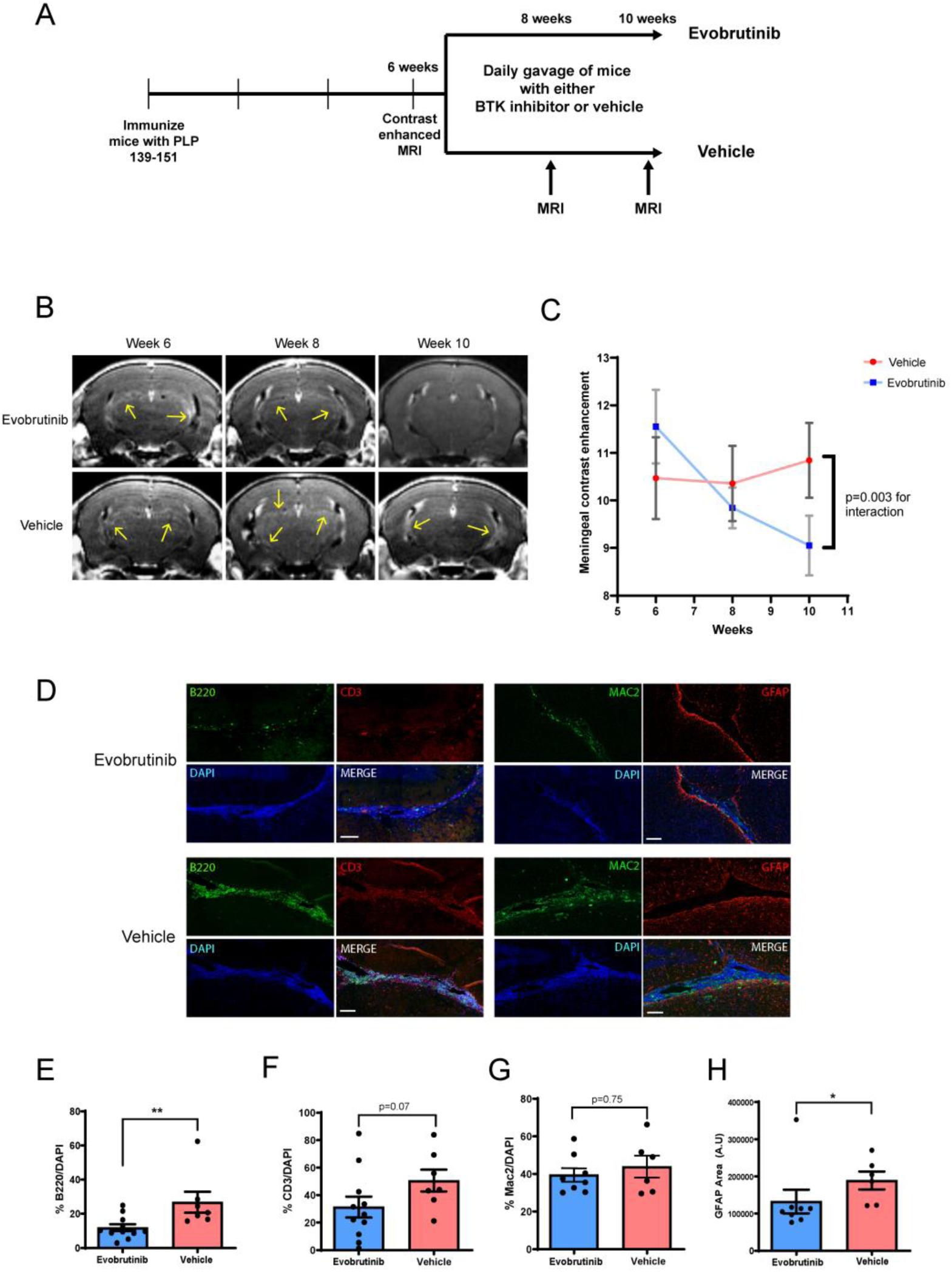
BTK inhibition reduced leptomeningeal inflammation detected on MRI and by histopathology. **(A)** Experimental design used to test effect of a BTK inhibitor (evobrutinib) on established leptomeningeal inflammation in EAE. Mice with evidence of leptomeningeal contrast enhancement were randomized to either vehicle or evobrutinib group and treated for 4 weeks. **(B)** We noted a reduction in contrast enhancement with evobrutinib treatment as compared to vehicle over the 4-week treatment period. This is quantified in **(C). (D and E)** On pathological evaluation at week 10, we noted reduction in the proportion of B cells within the meningeal infiltrate in the evobrutinib group compared to vehicle. A similar trend was noted for T cells but was not significant **(D and F)**. There was no significant difference in proportion of myeloid cells (Mac2+) **(D and G). (D and H)** We noted a reduction in GFAP+ cells in the cortex adjacent to the areas of meningeal inflammation in the evobrutinib group compared to vehicle. Scale bars in **D** are 100 µm. For **E–H** data is represented as mean ± SEM and *P* values are derived from a Mann-Whitney U test. *: *P* < 0.05, ** *P* < 0.005.

On pathological evaluation, we noted a significant reduction in the percentage of B cells within areas of meningeal inflammation (Fig. 7D and E). A similar trend was noted for T cells but this was not statistically significant (Fig. 7D and F). However, myeloid cell infiltrates in the meninges appeared to persist despite B cell depletion (Fig. 7D and G). We also noted a significant reduction in the astrocytosis in the cortex surrounding the areas of meningeal inflammation suggesting that changes in the meningeal infiltrate were impacting inflammation in the adjacent cortex (Fig. 7D and H).

These results demonstrate that BTK inhibitor treatment reduced meningeal inflammation by altering the immune cell composition of meningeal inflammatory infiltrates and resulted in changes in the adjacent cortex.

## Discussion

In this study we first demonstrated the ability of ultra-high field contrast-enhanced MRI, in SJL/J mice with EAE, to detect areas of meningeal contrast enhancement corresponding to meningeal inflammation. We then identified various pathological features of meningeal inflammation and their effects on the adjacent cortex that bear similarities to leptomeningeal inflammation in multiple sclerosis. Lastly, we demonstrated the utility of meningeal contrast enhancement as an outcome marker to assess the efficacy of treatment strategies targeting leptomeningeal inflammation. We note a potential therapeutic role of a BTK inhibitor – evobrutinib in ameliorating established meningeal inflammation in EAE, suggesting that this strategy warrants further investigation in human studies.

While imaging of meningeal inflammation has been described in multiple sclerosis both at 3 Tesla and at 7 Tesla magnet field strengths, only one other study has attempted to image this process in EAE (Pol *et al*., 2019). In their study Pol et al. noted potential MRI changes representing meningeal inflammation which peaked at day 10 post-immunization and then gradually declined over time. Unfortunately, the model chosen for that study does not develop persistent meningeal inflammation with significant B cell accumulation or the formation of tertiary lymphoid organs and hence it is unclear whether this approach would have produced results similar to those noted in our study if applied to the rr-EAE model (Magliozzi *et al*., 2004). We provide evidence that ultra-high field contrast enhanced MRI can not only detect areas of meningeal inflammation, but also track this process over time. This provides a potential surrogate marker of treatment response to help screen treatment modalities that impact meningeal inflammation.

We also characterized the meningeal infiltrate and noted results similar to prior studies, demonstrating that areas of meningeal inflammation had an abundance of B and T lymphocytes. In addition, we noted the presence of myeloid cells including follicular dendritic cells. We also identified the presence of PNAd+ vessel-like structures consistent with high-endothelial venules. Thus, we noted some features consistent with tertiary lymphoid organ formation as previously described in both EAE and multiple sclerosis. The unique aspect of our study was that we evaluated changes in the meningeal infiltrate over an extended time period in EAE and noted that the proportion of B cells within these areas continued to increase over the course of EAE – over 3-fold from week 2 to week 8–10 post-immunization. This is important when considering the timing of interventions that target this immune cell population.

In addition to examining the meningeal immune infiltrate, we also evaluated changes in the cortex adjacent to these areas. This is of interest since the major rationale to image and treat meningeal inflammation in multiple sclerosis is its potential relationship to increased cortical demyelination and neuronal loss. We identified the presence of increased axonal stress/damage, astrocytosis, microgliosis and demyelination in the cortex adjacent to areas of meningeal inflammation similar to those noted in multiple sclerosis. A recent study utilizing an adoptive transfer approach in SJL mice also demonstrated similar changes in the cortex adjacent to areas of meningeal inflammation (Ward *et al*., 2019).

The mechanism by which meningeal inflammation may contribute to the development of cortical changes is unknown. However, potential explanations include the production of antibodies or other soluble factors from B cells in the meninges, or the activation of macrophages/microglia or astrocytes in the cortex. We noted the presence of activated microglia/macrophages in the cortex and also noted C3+ and PSMB8+ astrocytes which could represent reactive neurotoxic astrocytes in the cortex adjacent to areas of meningeal inflammation. Additional studies are required to identify the precise mechanism mediating cortical pathology in this model.

Since multiple clinical trials targeting meningeal inflammation in multiple sclerosis with B cell depleting therapies have been unsuccessful, we decided to test an agent that targets both B lymphocytes and myeloid cells. BTK plays an important role in B cell development and in pro-inflammatory myeloid cell function (Gabhann *et al*., 2014; Haselmayer *et al*., 2019). We noted that over 4-weeks of treatment, evobrutinib reduced meningeal contrast enhancement compared to vehicle and on histopathology this corresponded to a decrease in B cells within the meningeal infiltrate. We also identified reduced astrocytosis in the adjacent cortex suggesting that the effects on the meningeal infiltrate led to reduced pathology in the surrounding cortex. Evobrutinib has been shown to reduce relapse rate in patients with relapsing remitting multiple sclerosis in a recent phase 2 trial with no significant serious adverse effects (Montalban *et al*., 2019). The results of the current study provide support for testing the effects of BTK inhibition in progressive multiple sclerosis and especially those with evidence of leptomeningeal inflammation.

In conclusion, we demonstrate the ability of ultra-high field contrast-enhanced MRI to detect and track meningeal inflammation in EAE and utilize this method to identify a novel potential therapeutic strategy to target meningeal inflammation in multiple sclerosis patients.

## Acknowledgements

We would like to thank Kazi Akther and Jaidi Xu for technical assistance with imaging of animals.

## Abbreviations

APP: amyloid precursor protein
BTK: Bruton’s tyrosine kinase
CFA: complete Freund’s adjuvant
EAE: experimental autoimmune encephalomyelitis
NGS: normal goat serum
PNAd: peripheral node addressin
RARE: rapid acquisition with refocused echoes
rr-EAE: relapsing remitting model of experimental autoimmune encephalomyelitis
TE: echo time
TR: repetition time

