## Supplementary Material for "Imaging meningeal inflammation in CNS autoimmunity identifies a therapeutic role for BTK inhibition"

**for**

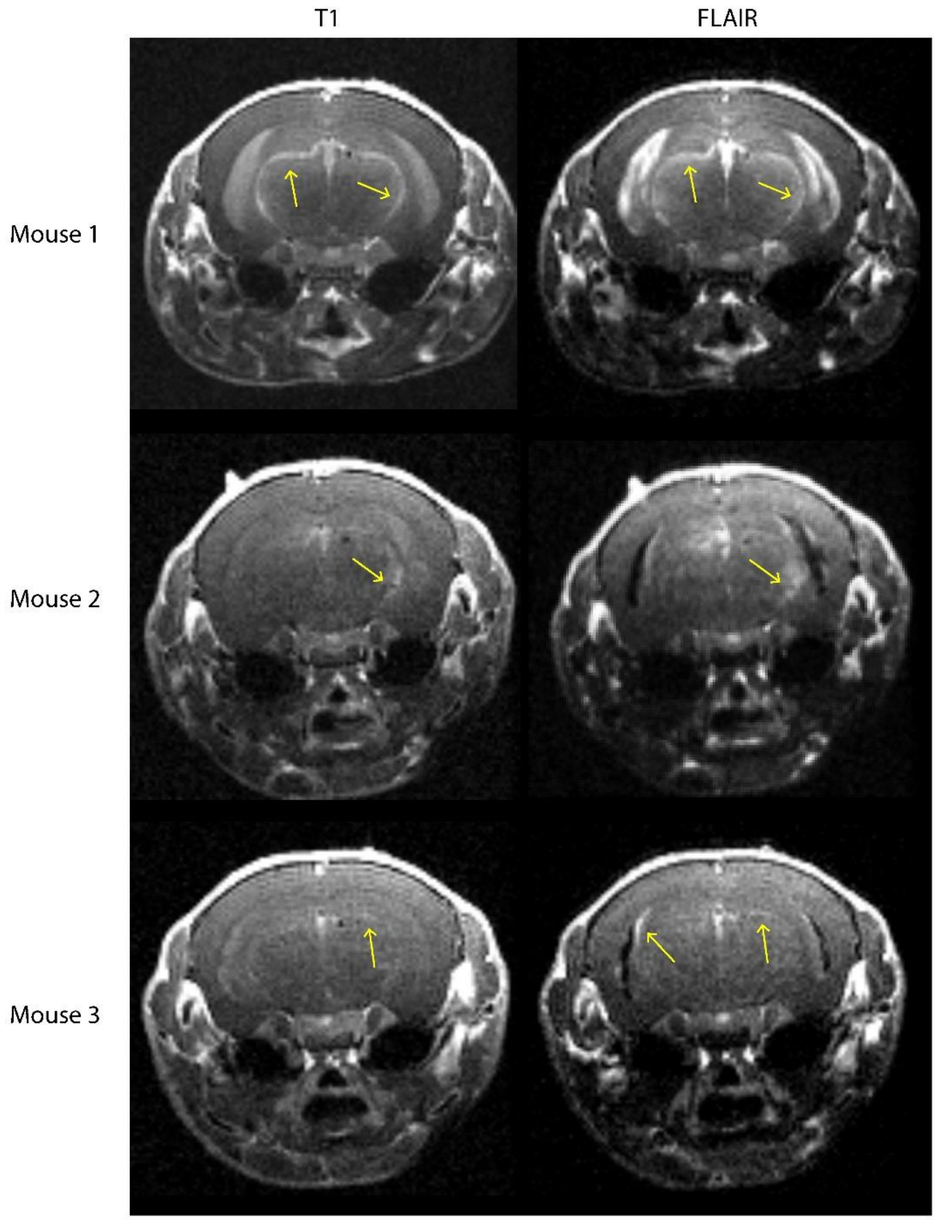


**Supplementary Figure 1. FLAIR imaging improves the sensitivity of imaging to detect areas of meningeal contrast enhancement in mice with EAE.** Examples of MRI images from mice with EAE using both T1 and FLAIR post-contrast sequences demonstrates that areas of meningeal contrast enhancement are more clearly demarcated and identified in the FLAIR post-contrast images.

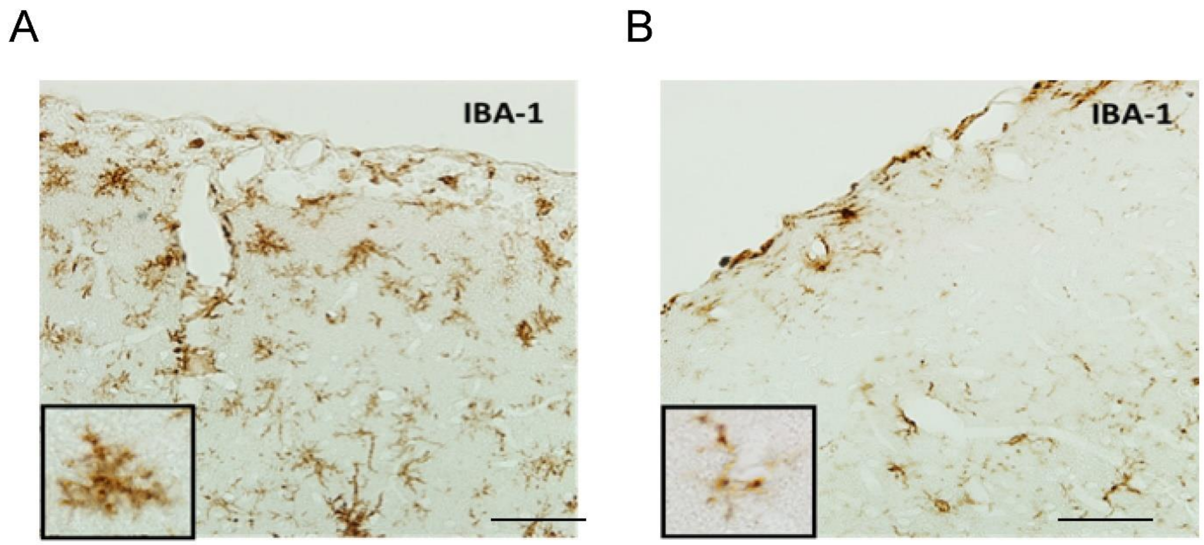


**Supplementary Figure 2. Microglia in the cortex adjacent to areas of meningeal inflammation demonstrate an activated phenotype.** (A) In the cortex adjacent to an area of meningeal inflammation Iba-1+ cells are increased in number and have a more ameboid appearance (A and inset) as compared to (B) the contralateral hemisphere where no meningeal inflammation was noted and the Iba-1+ cells have a resting appearance (B and inset).

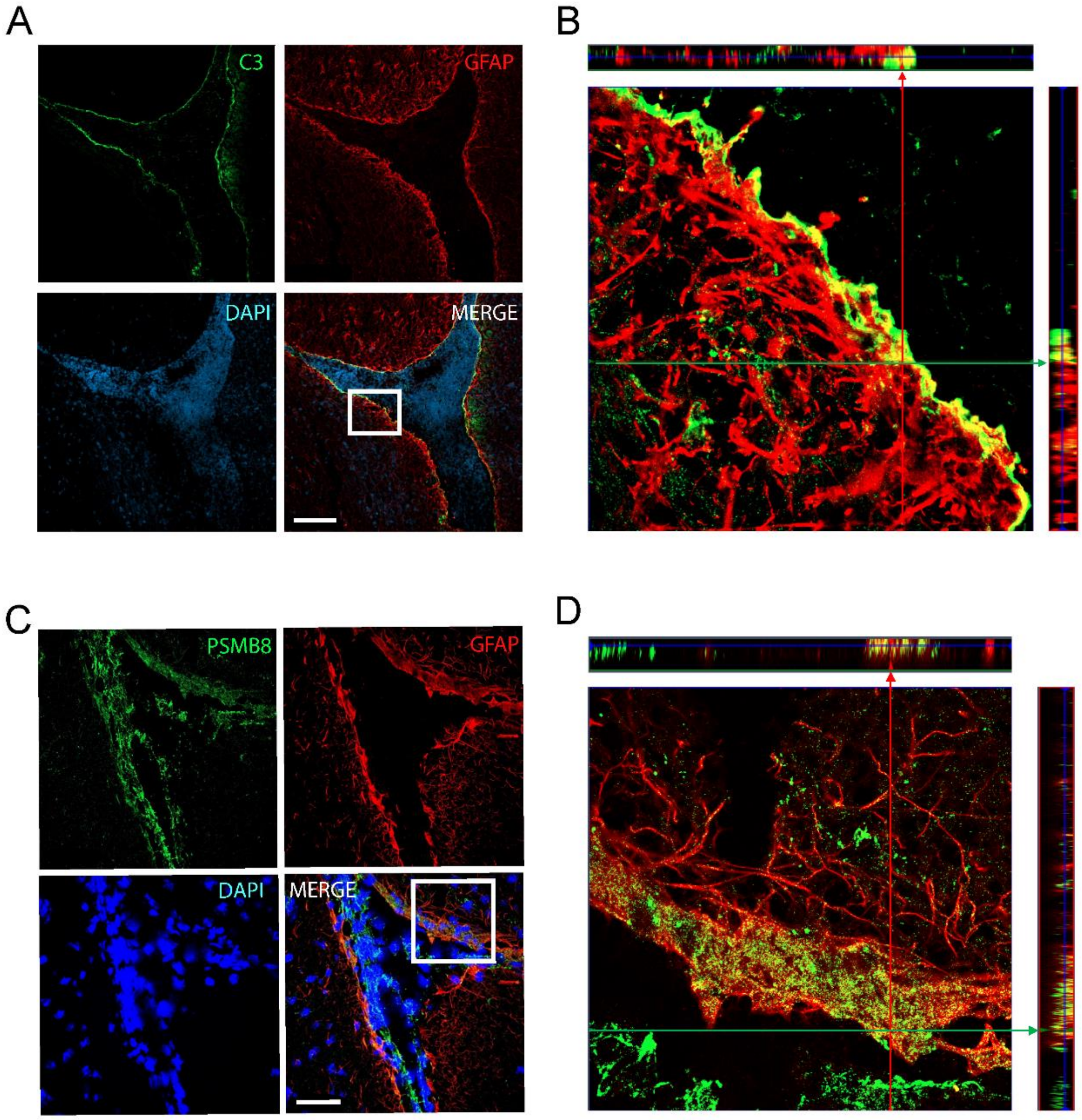


**Supplementary Figure 3. Inflammatory reactive astrocytes are present in cortex adjacent to leptomeningeal inflammation.** In mice with EAE demonstrating the presence of meningeal inflammation we performed double immunostaining for C3 and GFAP **(A)** and noted the presence of C3+ and GFAP+ astrocytes **(B)** in the cortex adjacent to meningeal inflammatory infiltrates. We also performed double immunostaining for PSMB8 and GFAP **(C)** and noted similar results with identification of PSMB8+ and GFAP+ astrocytes **(D)** in cortex adjacent to meningeal inflammation. Scale Bars: 100 µm for **(A)**; 50 µm for **(C)**.

**Supplementary Table 1 Antibodies utilized for immunohistochemistry of EAE tissues**

| **Antibody** | **Manufacturer** | **Clone** | **Isotype** | **Host** | **Concentration** | **Antigen retrieval** |
|---|---|---|---|---|---|---|
| APP | Millipore Sigma | Mono; 22C11 | IgG1 | Ms | 1:200 | Yes |
| B220 | Thermo Fisher Scientific | Mono; RA3-6B2 | IgG2a | Rat | 1:200 | Yes |
| CD3 | Agilent Dako | Poly | | Rb | 1:200 | No |
| CXCL13 | biorbyt | Poly | IgG | Rb | 1:500 | Yes |
| FDC-M1 | BD Biosciences | FDC-M1 | IgG2c | Ms | 1:50 | Yes |
| Fibrinogen | Agilent Dako | Poly | Ig | Rb | 1:100 | Yes |
| GFAP | Agilent Dako | Poly | IgG | Rb | 1:1000 | Yes |
| IBA-1 | Wako | Poly | | Rb | 1:300 | Yes |
| INOS | Santa Cruz Biotechnology | Mono; NOS2(C-11) | IgG1 | Ms | 1:1000 | No |
| LYVE-1 | abcam | Poly | IgG | Rb | 1:200 | Yes |
| Mac-2 | BioLegend | Mono; M3/38 | IgG2a | Rat | 1:200 | Yes |
| MBP | Novus Biologicals | Mono; R29.6 | IgG1 | Ms | 1:200 | Yes |
| PNAd | BioLegend | MECA-79 | IgM | Rat | 1:200 | Yes |
| SMI-32 | BioLegend | SMI32 | IgG1 | Ms | 1:500 | Yes |
| Black Gold | Millipore Sigma | Manufacturer's protocol | | | | |
| H&E | Vector Laboratories | Manufacturer's protocol | | | | |
| Luxol Fast Blue | IHC World | Manufacturer's protocol | | | | |
| **All antigen retrieval was performed with citrate buffer (pH6)* | | | | | | |

IgG = immunoglobulin
